# Sustainability in the laboratory: evaluating the reuseability of microtiter plates for PCR and fragment detection

**DOI:** 10.1101/2024.10.16.618673

**Authors:** Ane Liv Berthelsen, Anneke J. Paijmans, Jaume Forcada, Joseph Ivan Hoffman

## Abstract

Single-use plastics (SUPs) are indispensable in laboratory research, but their disposal contributes substantially to environmental pollution. Consequently, reusing common SUP items such as microtiter plates represents a promising strategy for improving laboratory sustainability. However, the key challenge lies in determining whether SUP reuse can be implemented without sacrificing data quality. To investigate this, we conducted a simple experiment to assess the impact of reusing microtiter plates on microsatellite genotyping accuracy. Plates previously used for PCR and fragment detection were cleaned opting for an environmentally friendly approach using regular soap and then reused. Our results indicate that, while reusing PCR plates significantly increases genotyping error rates due to residual DNA contamination, detection plates can be reused without compromising data quality. Our approach offers laboratories a practical and sustainable option for reducing SUP waste and costs while maintaining research integrity.

## 2. Introduction

Plastic waste has escalated into a global crisis. It is now highly improbable that any marine environments on Earth remain unaffected by plastic pollution [1]. Numerous policies and legislation aimed at managing plastic waste have been introduced at the national level (https://www.globalplasticlaws.org) and negotiations for a Global Plastics Treaty are currently taking place [2]. However, sustainable laboratory management remains largely voluntary [3] and in biological, agricultural and medical research, plastic waste was estimated at around 5.5 million tonnes annually a decade ago [4]. This figure has undoubtedly risen since, despite the potential of sustainable laboratory management to reduce energy consumption, lower carbon footprints and cut the financial costs of running research facilities [5,6].

Fortunately, sustainable research practices have been gaining increasing attention. Each year, on the third Tuesday of September, researchers collect their plastic waste and post on social media under the hashtag LabWasteDay to raise awareness about the waste generated daily in research laboratories [7,8]. Concurrently, initiatives such as the Laboratory Efficiency Assessment Framework (LEAF) [9] and My Green Lab [10] provide researchers with tools to evaluate the environmental impacts of their laboratories and offer practical recommendations for improvements [11]. Often, these are simple adjustments such as reducing freezer temperatures from -80 °C to -70 °C, which can lower energy consumption by nearly a third [12]. Furthermore, the 6R Concept, which was introduced in 2023, offers a framework for identifying sustainable solutions within existing protocols [13]. When applied to a neurobiology protocol, this framework achieved a 65% reduction in single-use plastic (SUP) waste [14].

SUPs have become a ubiquitous component of modern research laboratories globally. Their convenience, sterility, and affordability have made them indispensable for a diverse array of experiments and protocols. However, SUPs are often classified as non-recyclable due to bio-safety concerns or the risk of contamination [11,15]. As a result, much of the plastic waste produced by research laboratories is either incinerated or sent to landfills [7,14]. While glassware can sometimes provide a practical alternative, it is not a feasible option for many items like microtiter plates or pipette tips. Consequently, reusing common SUP items offers a promising avenue for enhancing laboratory sustainability [7].

Microtiter plates, otherwise known as microplates or microwell plates, are flat plates containing multiple wells that allow large numbers of samples to be processed simultaneously in small volumes. They are used for a wide variety of applications in analytical research and clinical diagnostics including immunoassays, colorimetric assays, tissue culturing and genetic screening [16]. The plastic polymers used in the manufacturing of microtiter plates vary depending on the specific application. For processes involving thermal cycling such as polymerase chain reaction (PCR), polypropylene is the most commonly used polymer because of its resistance to heat and chemicals. However, while these characteristics make polypropylene ideal for laboratory use, they also contribute to its prevalence in aquatic environments, where it is one of the most frequently detected polymers in plastic pollution [17].

PCR is a crucial step in the amplification of microsatellites, which are codominant genetic markers used widely in both academic and industrial settings [18]. Microsatellite loci are known for their high mutation rates, which give rise to high levels of allelic diversity [18]. These alleles are PCR amplified using oligonucleotide primers that bind to complementary sequences flanking the microsatellite. By incorporating fluorescent dyes into the PCR products, individual genotypes can be resolved through capillary electrophoresis and analysed using software packages such as GeneMarker (SoftGenetics, LLC, Pennsylvania, USA). This software implements the semi-automated calling of alleles by reference to marker panels containing information on expected allele sizes at each locus.

Unfortunately, even when sufficient amounts of high-quality DNA are available, microsatellites are susceptible to genotyping errors [19,20]. One of the primary sources of error is the mis-scoring of alleles, which can exacerbated by the presence of stutter bands (artefactual peaks resulting from slippage during PCR amplification) and signal intensity differences between alleles [20]. To guard against genotyping errors, it is essential to manually review the automated allele calls produced by fragment analysis software such as GeneMarker. Additionally, it is advisable for all genotype calls to be cross-checked by at least one independent, experienced observer. Several approaches can be used to detect microsatellite genotyping errors, with the gold standard being the blind re-genotyping of a subset of samples [20].

Our laboratory routinely uses microsatellites for population genetic studies of wild animal populations [21–25]. As part of a long-term project on Antarctic fur seals, we have genotyped around 15,000 individuals sampled over three decades at between nine (≤2009) and 39 (2010–present) microsatellite loci [26–28]. Our standard microsatellite genotyping protocol (see Methods for details) processes batches of 96 samples in two consecutive steps – PCR amplification and fragment detection – each using a different type of microtiter plate. In the first step, the 39 microsatellites are PCR amplified in five separate multiplexes, each on its own “PCR plate”. The second step involves diluting the amplified products, mixing them with a size standard, and transferring them to “detection plates” for analysis on an automated capillary sequencer. This procedure requires a total of ten microtiter plates for each batch of 96 samples.

To reduce SUP consumption in our laboratory, we performed a simple experiment to evaluate whether the PCR and / or detection plates could be reused without sacrificing data quality. We established four treatment groups within our experimental workflow: (i) “standard protocol”, (ii) “internal control”, (iii) “reused PCR plate” and (iv) “reused detection plate” (Figure 1). The standard protocol, as previously described, used new PCR and detection plates (i.e. plates that had never been used before) and served as a reference for comparison with the other treatment groups. The internal control was identical to the standard protocol, also using new plates. By comparing the genotypes obtained from the standard protocol and the internal control, we could determine the “baseline” genotyping error rate.

**Figure 1:**
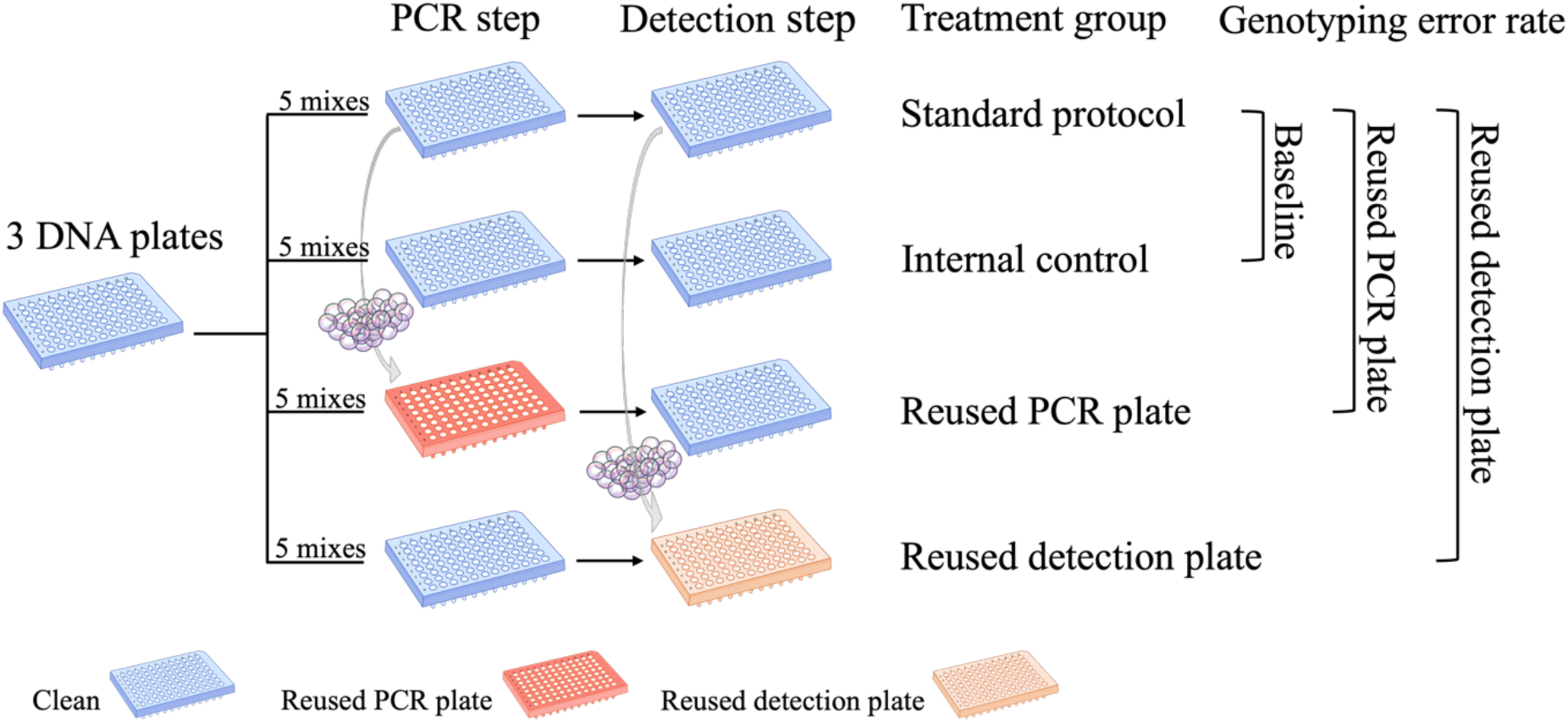
Overview of our experimental setup, which included four treatment groups. The “standard protocol” treatment group used new PCR and detection plates and served as a reference to quantify genotyping error rates for the other treatment groups. The “internal control” treatment group also used new PCR and detection plates and was used to estimate the baseline rate of genotyping error. The PCR plates and detection plates from the standard protocol treatment group were cleaned and reassigned to the “reused PCR plate” and “reused detection plate” treatment groups respectively. A total of three plates of DNA samples (n = 288) were genotyped at five multiplexes for each treatment group. The plates are colour coded according to the legend. Original artwork by A.L.B..

As described in the Methods, we then cleaned the PCR and detection plates used in the standard protocol and reassigned them for use in the third and fourth treatment groups. In the third treatment group, reused PCR plates were employed for the PCR step, while new detection plates were used for fragment detection. In the fourth treatment group, new PCR plates were used for the PCR step, while reused detection plates were employed for fragment detection. By comparing the genotypes obtained from the standard protocol and the third treatment group, we evaluated the impact of reusing PCR plates on the genotyping error rate. Similarly, by comparing the genotypes obtained from the standard protocol and the fourth treatment group, we assessed the impact of reusing detection plates on genotyping accuracy.

We processed a total of 288 samples across three 96-well microtiter plates, following the experimental workflow outlined above. To evaluate the effects of reusing microtiter plates at different stages of the protocol, we implemented a formal Bayesian analysis of genotyping error rates. We hypothesised that (i) reusing microtiter plates should be feasible, in at least some circumstances, without significantly compromising data quality; however (ii) the high sensitivity of PCR to trace amounts of DNA might introduce a risk of cross-contamination when reusing PCR plates, potentially increasing the genotyping error rate. Conversely, (iii) we anticipated that reusing detection plates would likely have a minimal impact on the genotyping error rate, as the capillary sequencer measures all signals, but only the strongest signals are scored.

## 3. Materials and Methods

### Tissue sampling and DNA extraction

Tissue samples were collected from 288 Antarctic fur seals from an intensively studied breeding population at Bird Island, South Georgia (54°00024.800 S, 38°03004.100 W) during the austral summers of 2006–2007, 2015– 2016, and 2020–2021. The seals were captured and restrained following protocols that have been established over more than 40 consecutive years of the long-term monitoring and survey program of the British Antarctic Survey (BAS). Pups were captured with a noosing pole on the day of birth and sampled from the umbilicus using piglet ear notching plyers. Each sample was stored individually in 20% dimethyl sulphoxide (DMSO) saturated with salt at −20 °C. Total genomic DNA was later extracted using an adapted chloroform-isoamylalcohol protocol [29].

### Standard microsatellite genotyping protocol

All samples were genotyped following our standard protocol, as detailed by Paijmans et al. [30]. In brief, 39 microsatellite loci were PCR amplified in five separate multiplex reactions using a Type It Kit (Qiagen). For this step, we used ultra-thin walled, non-skirted PCR plates (PCR trays ROTILABO® 96 well, Standard, half frame, Roth Selection, Karlsruhe, Germany). Each plate contained 96 samples including three positive controls to facilitate the standardisation of microsatellite allele calling across plates. The PCR program included an initial denaturation step of 5 minutes at 94°C, followed by 28 cycles of 30 seconds at 94°C, 90 seconds at the annealing temperature (Ta°C) specified for each multiplex reaction, and 30 seconds at 72°C, with a final extension of 30 minutes at 60°C. The fluorescently labelled PCR products were transferred to hard-shell, fully-skirted detection plates (Fisherbrand™ 96-Well Semi-Skirted PCR Plates, Thermo Fisher Scientific, Waltham, MA, USA) before resolving them by electrophoresis on an ABI 3730xl capillary sequencer (Applied Biosystems, Thermo Fisher Scientific, Waltham, MA, USA). Allele sizes were automatically scored using GeneMarker v. 2.6.2 (SoftGenetics, LLC., State College, PA, USA) and the traces were manually inspected by two independent observers (A.L.B and J.I.H., or A.P and J.I.H.), with corrections being made where necessary to maximise genotype quality.

### Experimental design

As outlined in the introduction and illustrated in Figure 1, our experiment included four treatment groups. The first (standard protocol) and the second (internal control) treatment group followed the previously described workflow, both using new PCR and detection plates. We opted for a gentle, environmentally friendly approach to clean the plates by using regular soap as described here: each plate was rinsed with distilled water and emptied ten times, before submerging it in soapy water for two hours. After soaking, the plates were again rinsed and emptied ten times before being left overnight on a paper towel to dry. The cleaned PCR and detection plates were subsequently reassigned to treatment groups three and four respectively. We retained information about the samples that were originally processed on each plate and ensured that no plate was reused for the same samples originally processed on it.

### Evaluation of genotyping errors

Genotyping error rates were calculated based on discrepancies between the genotypes obtained from the standard protocol (reference treatment group) and those obtained from the three other treatment groups (internal control, reused PCR plate and reused detection plate). For each single-locus genotype, a binomial variable “mismatch” (0 = match, 1 = mismatch) was computed. A “match” indicates complete agreement between the genotypes from the standard protocol and the tested treatment group, while a “mismatch” indicates a discrepancy at one or both alleles. Following Hoffman and Amos [20], we then calculated the error rate per reaction as the number of mismatching single-locus genotypes divided by the total number of genotypes compared.

### Statistical analysis

We fitted a binomial Bayesian logistic mixed effect model to evaluate the effects of reusing PCR and detection plates on genotyping error rates. The response variable was “mismatch” and treatment group was included as a three-level fixed effect explanatory variable, with the internal control treatment group set as the reference (intercept) category. To account for heterogeneity arising from the use of different samples, DNA plates, multiplexes and loci, these variables were included as random effects in the model as follows:

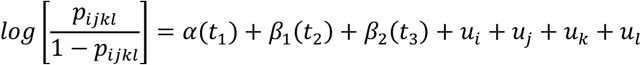

Where pijkl represents the probability that the binary response variable mismatch is equal to one (mismatch observed) for an observation within the levels of the random effect variables sample ID (indexed by *i*), DNA plate (indexed by *j*), multiplexed reaction (indexed by *k*) and loci (indexed by *l*). The internal control treatment group was set as reference category (α(t1)), which the other treatment groups were compared against. This analysis was performed using the *brms* package version 2.20.4 [31], with three independent Markov chains being run for 100,000 iterations after a burn-in of 30,000 iterations with a thinning interval of 70. The trace plots were visually inspected and model diagnostics such as R hat statistics and autocorrelation were generated using *brms*. All data analyses were implemented in R version 4.2.1 [32] with Rstudio version 2023.09.1+494 [33].

## 4. Results

Multilocus genotypes were successfully generated for all four treatment groups for a total of 281 samples (97.6%). The corresponding genotyping error rates, calculated per reaction by reference to the standard protocol, are shown in Table 1. The internal control and reused detection plate treatment groups exhibited similarly low genotyping error rates, at 0.005 and 0.004 per reaction respectively. However, the genotyping error rate for the reused PCR plate treatment group was approximately ten times higher, at 0.034 per reaction.

**Table 1:**
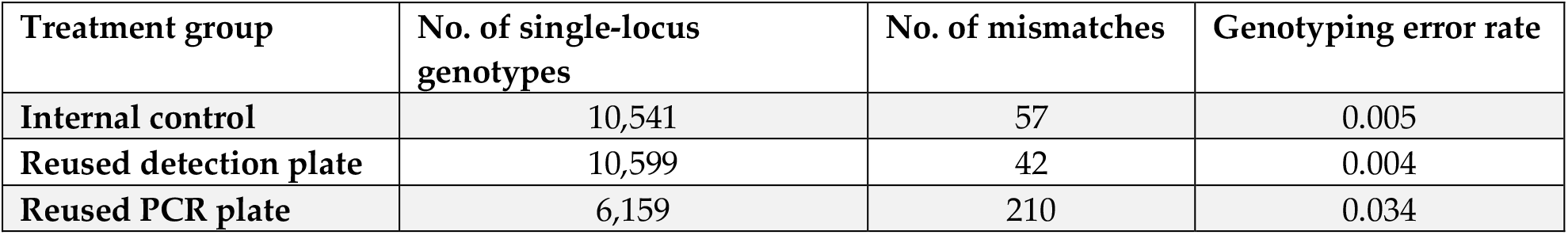
Per-reaction genotyping error rates for the internal control, reused detection plate and reused PCR plate treatment groups, calculated relative to the standard protocol treatment group. Due to variable amounts of missing data, the number of single-locus genotypes differs among the treatment groups.

To formally analyse variation in genotyping error rates among the treatment groups while accounting for heterogeneity among different samples, DNA plates, multiplexes and loci, we implemented a Bayesian logistic mixed effects model as described in the Methods. The model accounted for nearly 12% of the total variation in genotyping errors. The posterior distributions of the standardized beta coefficients of the genotyping error rates of the internal control and the reused detection plate treatment groups were similar and their 95% confidence intervals (CIs) overlapped (Figure 2, Table 2). This indicates that reusing detection plates did not lead to a measurable increase in the genotyping error rate compared to the internal control. By contrast, the posterior distribution of the standardized beta coefficients of the genotyping error rate of the reused PCR treatment group was considerably higher than that of the internal control, with the 95% CIs of the two treatment groups not overlapping (Figure 2, Table 2). This indicates that reusing PCR plates resulted in a significantly higher genotyping error rate.

**Table 2:**
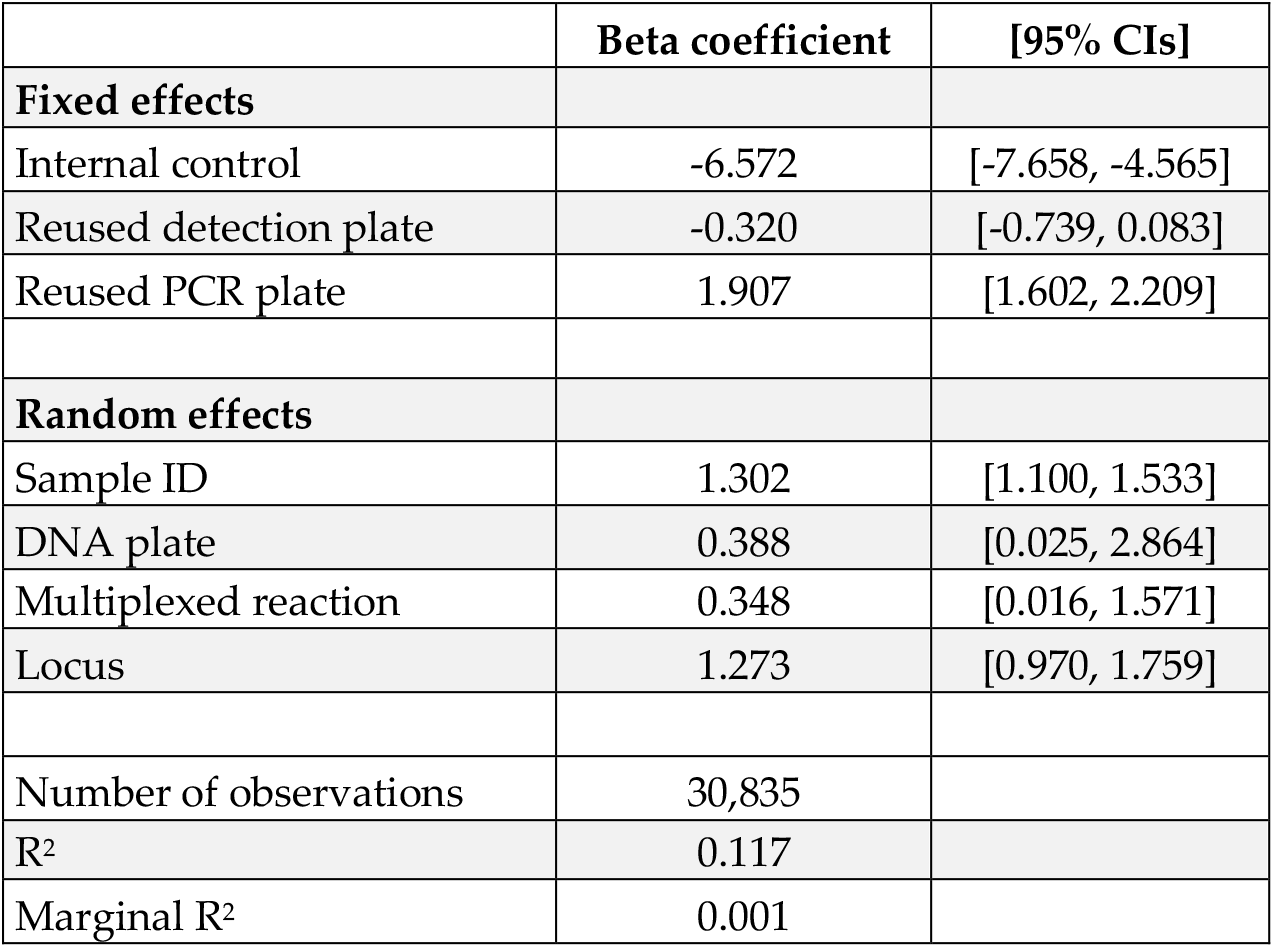
Standardized point beta coefficients and 95% confidence intervals (in square parentheses) from the Bayesian logistic mixed effect model testing for the effects of the three-level categorical fixed effect “treatment” on genotype errors while controlling for the random effects sample ID, DNA plate, multiplexed reaction and loci. Reused PCR plate and Reused detection plate estimates are compared against to the internal control treatment group. Shown also are the R2 (the proportion of the total variance explained by the model) and marginal R2 (the proportion of the total variance explained by the fixed effects alone).

**Figure 2:**
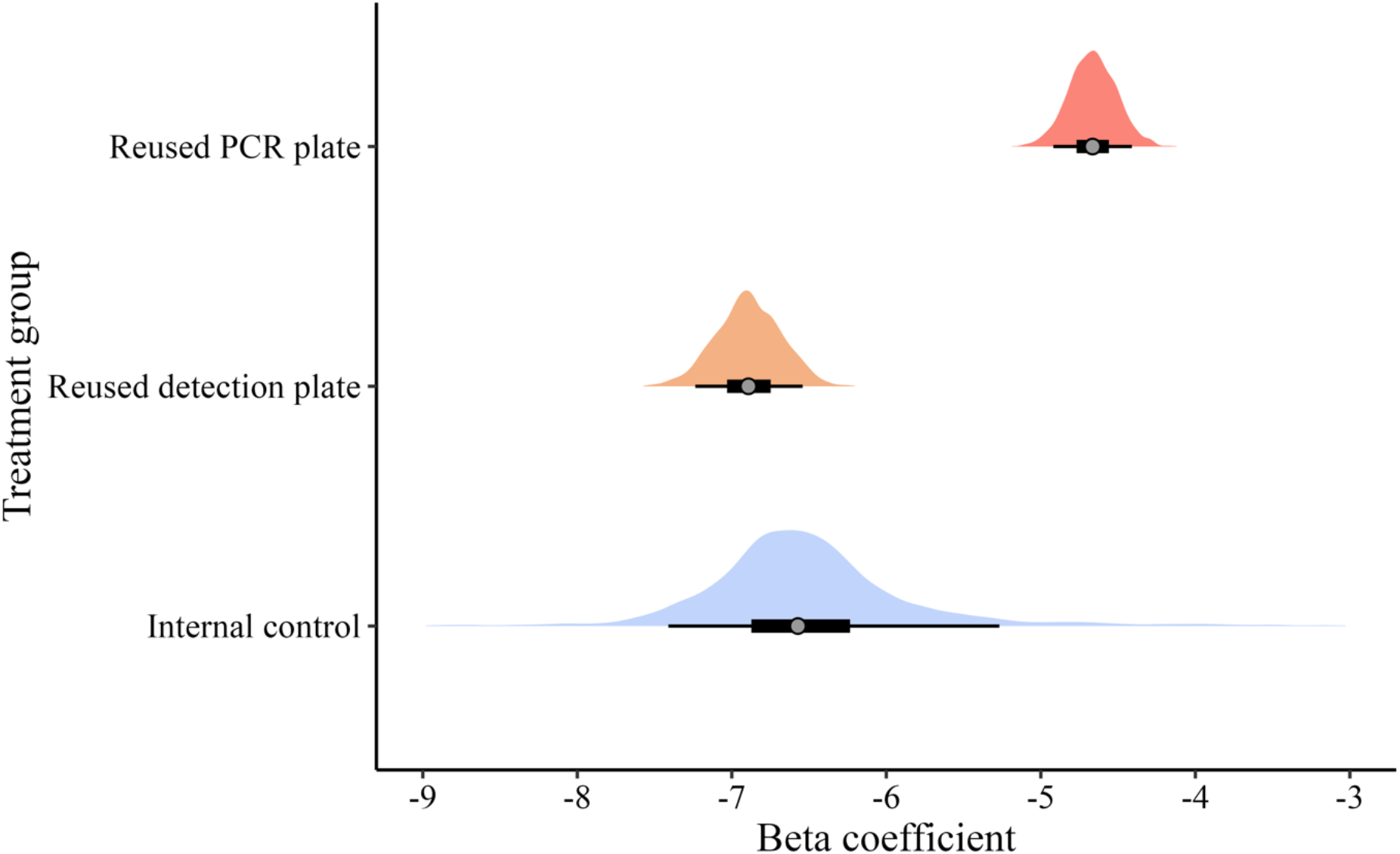
Posterior distributions of the standardized beta coefficients of the internal control (blue), reused detection plate (orange) and reused PCR plate (red) treatment groups on genotyping errors. The grey points represent the mean posterior estimates, the thick black lines represent the 50% confidence intervals, and the thin black lines represent the 90% confidence intervals.

## 5. Discussion

We conducted a simple experiment to investigate the potential for reusing microtiter plates in our laboratory. We found that genotyping error rates for both the internal control and reused detection plate treatment groups were similar in magnitude to genotyping error rates observed in previous microsatellite-based studies from our group, which range from 0.0013–0.0074 per reaction [20,25,34]. Importantly, these error rates fall well below the one percent threshold generally considered acceptable for microsatellite genotyping [35]. This indicates that detection plates can be reused in our fur seal genotyping protocol without compromising data quality. Nevertheless, despite this promising result, we recommend proceeding with caution. As we move forward with detection plate reuse in our laboratory, it will be important to maintain strict quality control by regularly monitoring genotyping error rates. This measure will help to ensure that any potential issues are promptly identified and addressed, safeguarding the integrity of our research.

The genotyping error rate for the reused PCR plate treatment group was nearly ten times higher than the background rate, exceeding the one percent threshold by a factor of three. This indicates that reusing PCR plates led to an unacceptable loss of data quality. This supports our original hypothesis that the high sensitivity of PCR to residual DNA may elevate genotyping error rates when reusing these plates. In contrast, the fragment detection step is less sensitive to residual DNA, as only the strongest signals are scored, and florescent signals are degraded by our cleaning protocol. However, we employed a relatively gentle cleaning method, which appears to have been insufficient to remove all traces of DNA. Alternative approaches, such as treating plates with DNAse or 10% bleach, could be more effective. Other options include UV irradiation or autoclaving, although these methods risk damaging the plates, unless care is taken to ensure the plates can withstand the treatment, e.g. autoclavable plates (https://www.carlroth.com/de/de/pcr-platten/pcr-platten-96-well/p/eyt4.1). Testing these alternatives would be worthwhile, but their effectiveness is uncertain as polypropylene is known to adsorb DNA [36]. This inherent limitation of polypropylene may limit the success of cleaning methods, although using low attachment plates made of polyallomer or low-binding polypropylene might help to mitigate this issue.

Unexpectedly, the reused PCR plate treatment group also exhibited a higher percentage of missing data (13.4%) compared to both the internal control (3.0%) and the reused detection plate (2.3%) treatment groups. To investigate this further, we revisited the original GeneMarker projects and classified the missing data into three categories: (i) failed PCR reactions, characterized by weak or absent PCR products; (ii) uninterpretable PCR reactions, which produced high-intensity peaks that did not resemble microsatellite alleles; and (iii) probable contamination cases, where the genotypes could not be scored due to the presence of more than two alleles. These categories accounted for 48.7%, 24.5% and 26.9% of the failed reactions respectively. This indicates that PCR amplification was generally less reliable on the reused PCR plates, with potential cases of contamination only explaining about one quarter of the missing data.

While cleaning PCR plates may pose challenges, reusing detection plates in our microsatellite genotyping protocol offers significant benefits in terms of reducing SUP consumption and costs. The extent of these savings will depend on how many times each detection plate can be reused, which is yet to be determined. However, even if each detection plate were reused just once, SUP consumption would decrease by around 100g, leading to a saving of around 33.20 EUR for the genotyping of each batch of 96 Antarctic fur seals. If the detection plates could be reused multiple times, these savings would be even greater. Currently, we estimate that our department uses around 500 detection plates per year. Reducing this number by 50% would save 5 kilograms of SUP and approximately 1,660.00 EUR in consumables annually. Although these savings may seem modest at a first glance, every step contributes to reducing our environmental footprint, and the cumulative benefits of implementing this simple measure will grow over time.

It is important to note that our findings are specific to our experimental protocol and may not be directly applicable to other laboratories and contexts without further validation. Additionally, we recognize that reusing or recycling microtiter plates and other laboratory equipment may not always be appropriate, particularly in scenarios where precision is paramount, such as in medical diagnostics. However, in less critical settings, we see considerable potential for reusing microtiter plates. We believe that our findings can also serve as a foundation for further research into the safe and effective reuse of laboratory materials across diverse research environments, and so we encourage our colleagues to explore the potential for reusing SUPs in their laboratories.

## 6. Conclusion

Our findings demonstrate that a common SUP item – microtiter plates – can be reused under certain conditions without any notable decline in data quality. This approach offers dual benefits: reducing plastic waste while also lowering costs, making it especially appealing for research groups or institutions operating on limited budgets. However, we emphasize the importance of continuous quality control to ensure consistently high data standards. More broadly, other SUPs in laboratory settings might also be suitable for reuse, offering further opportunities for waste reduction across various scientific disciplines.

## Acknowledgments

The authors would like to thank Janina Mercedes Grabowski and Prisca Viehöver for their valuable contributions to the laboratory work. This research was funded by the Deutsche Forschungsgemeinschaft (DFG, German Research Foundation) Collaborative Research Centre Transregio 212 “A Novel Synthesis of Individualisation across Behaviour, Ecology and Evolution: Niche Choice, Niche Conformance, Niche Construction (NC3)” (SFB TRR 212, project numbers 316099922 & 396774617) and by the DFG special priority programme “Antarctic Research with Comparative Investigations in Arctic Ice Areas” (SPP 1158, project number 424119118). Support for the article processing charge was granted by the DFG and the Open Access Publication Fund of Bielefeld University.

## Ethical Statement

Antarctic fur seal samples were collected at Bird Island, South Georgia under permits from the Government of South Georgia and the South Sandwich Islands (GSGSSI, Wildlife and Protected Areas Ordinance (2011), RAP permit numbers 2018/024 and 2019/032) by the British Antarctic Survey as part of the Polar Science for Planet Earth program. First, the samples were imported under permits from the Department for Environment, Food, and Rural Affairs (Animal Health Act, import license number ITIMP18.1397) and from the Convention on International Trade in Endangered Species of Wild Fauna and Flora (import numbers 578938/01-15 and 590196/01-18) into the United Kingdom. Second, the samples were exported to Germany under permits issued by the GSGSSI and the UK Department for Environment, Food and Rural Affairs, under European Communities Act 1972. British Antarctic Survey Animal Welfare and Ethics Review Body (reference no. PEA6, AWERB applications 2018/1050 and 2019/1058) approved all procedures, which were carried out in agreement with relevant guidelines and regulations, and reported in accordance with ARRIVE guidelines where applicable.

## Funding Statement

Ane Liv Berthelsen and Joseph Ivan Hoffman: SFB TRR 212 (NC^3^) (Project Numbers 316099922 & 396774617)

Anneke J. Paijmans and Joseph Ivan Hoffman: Deutsche Forschungsgemeinschaft (DFG, German Research Foundation) priority programme “Antarctic Research with Comparative Investigations in Arctic Ice Areas” SPP 1158 (project number 424119118)

Jaume Forcada: the core science programme of the British Antarctic Survey, Polar Science For a Sustainable Planet, NERC-UKRI (Natural Environment Research Council, United Kingdom UKRI).

## Data Accessibility

The script needed to reproduce the presented analysis, figure and tables has been provided as a R Quarto PDF file. It can additionally be accessed via Github https://github.com/AneLivB/SGP. The data are available via Zenodo [37].

## Competing Interests

The authors declare no competing interests.

## Authors’ Contributions

J.I.H. and A.L.B conceived the study and J.F. collected the samples. A.L.B and A.P. performed the genotyping and scoring of the microsatellite data, which was double checked by J.I.H. The manuscript was drafted by A.L.B. and J.I.H. All of the authors approved the final version.

## Notes

https://zenodo.org/records/13913891

https://github.com/AneLivB/SGP

### Competing Interest Statement

The authors have declared no competing interest.

