## Supplemental information for "Sustainability in the laboratory: evaluating the reuseability of microtiter plates for PCR and fragment detection"

Script

Ane Liv Berthelsen

2024-10-11

**Session information** **17**

This file contains the R code used for the manuscript "*Sustainability in the laboratory: evaluating the reuseability of microtiter plates for PCR and fragment detection*" by Ane Liv Berthelsen, Anneke J. Paijmans, Jaume Forcada and Joseph I. Hoffman. The data files can be downloaded via Zenodo, https://doi.org/10.5281/zenodo.13913891 and additional scripts are available on Github, https://github.com/AneLivB/SGP.

#### Aims

We set out to explore the reuseability of 96-microtiter plates with the aim to test the following hypothesis: (i) reusing microtiter plates should be feasible, in at least some circumstances, without significantly compromising data quality; however (ii) the high sensitivity of PCR to trace amounts of DNA might introduce a risk of cross-contamination when reusing PCR plates, potentially increasing the genotyping error rate. Conversely, (iii) we anticipated that reusing detection plates would likely have a minimal impact on the genotyping error rate, as the capillary sequencer measures all signals, but only the strongest signals are scored.

### Materials and Methods

We designed an experimental setup to assess the re-usability of 96-micro well plates within the context of microsatellite genotyping. The setup contained 4 treatments: standard procedure, internal control, re-used PCR plate and re-used detection plate (See Figure 1). Each Antarctic fur seal tissue sample was extracted using an adapted chloroform-isoamylalcohol protocol and genotyped following our standard protocol as detailed in Paijmans et al. 2024 (https://doi.org/10.1038/s41598-024-62290-x).

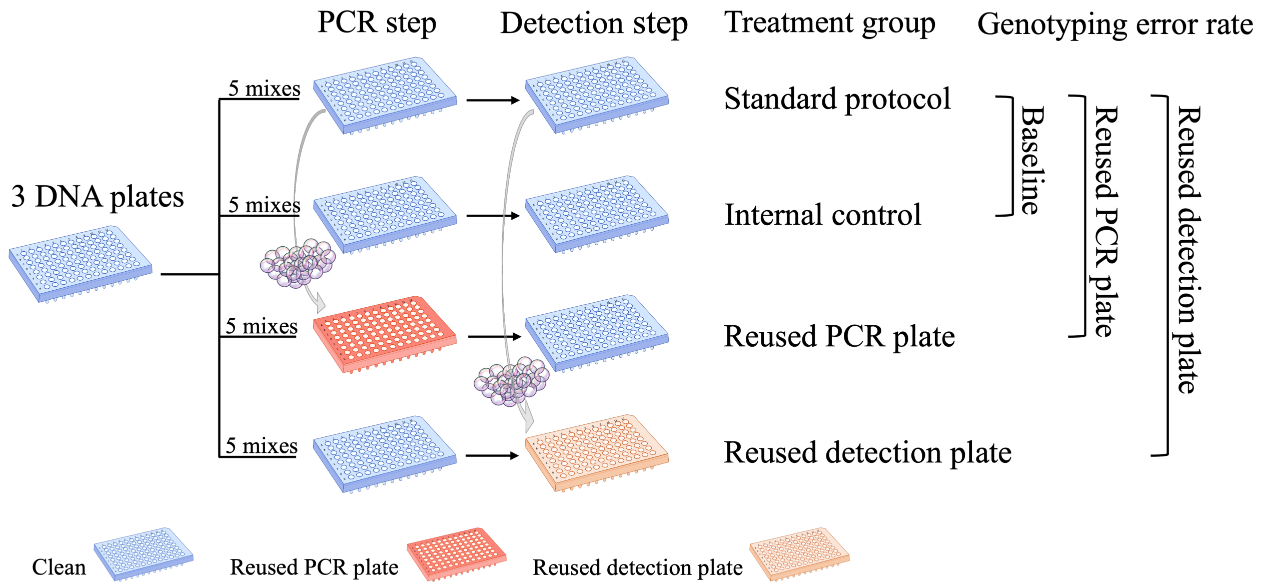

Figure 1: Schematic overview

The first (standard protocol) and the second (internal control) treatment group followed the standard protocol. Afterward, the plates from the standard protocol were cleaned. The cleaned PCR and detection plates were reassigned to treatment groups three and four respectively. We retained information about the samples that were originally processed on each plate and ensured that no plate was reused for the same samples originally processed on it. The genotyping error rate for each treatment was calculated by comparing the scored genotype to the corresponding genotype in the standard protocol treatment.

##### PCR Program

Table 1: PCR program

| Stage | No. of cycles | Temperature (°C) | Duration | Process |
|---|---|---|---|---|
| 1 | 1 | 94 | 5 minutes | Heat up |
| 2 | 28 | 94 | 30 seconds | Denaturation |
| | | 60/53* | 90 seconds | Annealing |
| | | 72 | 30 seconds | Extension |
| 3 | 1 | 60/53* | 30 minutes | Annealing |
| 4 | 1 | 10 | Hold | Cool down |

*Note: Annealing temperatures are mix-specific.*

### Script

#### Packages

```
#here::here()
invisible(pacman::p_load(dplyr, tidyverse, here, stringi, png, knitr, rstatix, kableExtra,
                         rstan, brms, bayesplot, ggridges, gt, tibble, webshot2))
#here: file referencing within project
#png to save png files
#gt for model output table
#bayesplot and ggridges for plot with beta distribution
#tibble for adding extra rows to table
#webshot2 to make table into png
#knitr and kableExtra for tables
#rstan and brms for models and model diagnostics
```

#### Data

The data for this project are the raw sequencing reads from the ABI 3730xl capillary sequencer. Each files is identified with the following name structure: RackX_mixY_Z. Where X gives the rack number, Y the mix and Z the treatment (1 = standard procedure, 2 = internal control, 3 = Re-used detection plate and 4 = Re-used PCR plate). The 0_Raw_data_processing.qmd script processes this data into .csv files containing the scored allele information for each DNA plate for all mixes, which is loaded in here.

```
#Load in the processed data files
Processed_data_files <- list.files(path = working_dir, pattern = "*.csv", full.names = T)
list2env(
  lapply(setNames(Processed_data_files,
                  make.names(gsub("Data/Working_data/Processed_data/|.csv", "",
                                  Processed_data_files))), read.csv),
  envir = .GlobalEnv)
R156_R3$mix3.ZcwDz301_a <- as.integer(R156_R3$mix3.ZcwDz301_a)
R156_R3$mix3.ZcwDh4.7_a <- as.integer(R156_R3$mix3.ZcwDh4.7_a)
```

##### Prepare the datasets

For each sample we noted whether the genotype matched with the ‘standard procedure’ per loci and for all treatments whether the genotype was scored or not. This yielded two variables referred to as ‘mismatch’ and ‘missing’ respectively. The binary mismatch variable (0 = match, 1 = mismatch) was used as the outcome variable for the model.

```
# #The dataset created from this function is called 'Analysis.data'
# #Analysis.data is the dataset used for the model.
# #The dataset can be loaded directly, so it's not necessary to run this function.

# Mismatch_loci <- function(DF1, DF2) {
#   Loci <- rbind(as.data.frame((colnames(DF1[3:80]))))
#   colnames(Loci) <- "Loci"
#   Loci$Loci <- gsub("_a", "", Loci$Loci)
#   Loci$Loci <- gsub("_b", "", Loci$Loci)
#   Loci <- unique(Loci$Loci)
#   Loci <- as.data.frame(Loci)
#   Loci$Mix <- substring(Loci$Loci, 1, 4)
#   Loci$Loci <- gsub("mix\\d{1}.", "", Loci$Loci)
#   Loci$Rack <- substring(deparse(substitute(DF1)), 1, 4)
#
```

```
#   #To recognize treatment, simplify the 'name' of the DF to the round number (R*)
#   Round <- gsub("R\\d{3}|_", "", deparse(substitute(DF2)))
#   if (Round == "R2"){
#     Treatment <- "Internal control"
#   } else if (Round == "R3") {
#     Treatment <- "Re-used detection plate"
#   } else {
#     Treatment <- "Re-used PCR plate"
#   }
#   Loci$Treatment <- Treatment
#
#   #To create the binomial match/no match column, first a base is created
#   Base <- DF1[1:2]
#   colnames(Base) <- c("Well", "ID")
#   Base$Row <- substring(Base$ID, 1, 1)
#
#   Loci <- cross_join(Loci, Base)
#   Loci$pair <- paste(Loci$ID, Loci$Loci)
#
#   #mismatches are identified and the row+column is merged
#   Mismatch <- as.data.frame(which(DF1[3:80] != DF2[3:80], arr.ind = TRUE))
#   Mismatch$row <- DF1$mix1.ID[Mismatch[,1]]
#   Mismatch$col <- Mismatch[,2]+2
#   Mismatch$col <- colnames(DF1[Mismatch[,2]])
#   Mismatch$col <- gsub("mix\\d{1}.|_a.\\d+", "", Mismatch$col)
#   Mismatch$col <- gsub("_b.\\d+", "", Mismatch$col)
#   Mismatch$col <- gsub("_a", "", Mismatch$col)
#   Mismatch$col <- gsub("_b", "", Mismatch$col)
#   Mismatch <- Mismatch[!duplicated(Mismatch[1:2]),]
#   Mismatch$pair <- paste(Mismatch$row, Mismatch$col)
#
#   #Match column created by matching loci pair with mismatch row+column pair
#   Loci$Match <- ifelse(Loci$pair %in% Mismatch$pair, 0, 1)
#
#   #Add column with NA values
#   Detect_NA <- as.data.frame(which(is.na(DF2[3:80]), arr.ind = TRUE))
#   Detect_NA$row <- DF1$mix1.ID[Detect_NA[,1]]
#   Detect_NA$col <- Detect_NA[,2]+2
#   Detect_NA$col <- colnames(DF1[Detect_NA[,2]])
#   Detect_NA$col <- gsub("mix\\d{1}.|_a.\\d+", "", Detect_NA$col)
#   Detect_NA$col <- gsub("_b.\\d+", "", Detect_NA$col)
#   Detect_NA$col <- gsub("_a", "", Detect_NA$col)
#   Detect_NA$col <- gsub("_b", "", Detect_NA$col)
#   Detect_NA <- Detect_NA[!duplicated(Detect_NA[1:2]),]
#   Detect_NA$pair <- paste(Detect_NA$row, Detect_NA$col)
#
#   #Match and Missing data coloum created by matching loci pair with mismatch row+column pair
#   #If a mismatch or missing data is observed, 0, otherwise 1
#   Loci$Mismatch <- ifelse(Loci$pair %in% Mismatch$pair, 1, 0)
#   Loci$Missing <- ifelse(Loci$pair %in% Detect_NA$pair, 0, 1)
#   #The unscored loci from the missing category, cannot be assessed for genotyping errors!
#   Loci$Mismatch <- ifelse(Loci$Missing == 0, NA, Loci$Mismatch)
#
#   Loci <- Loci[,-grep("pair",colnames(Loci))]
#   Loci$ID <- substring(Loci$ID, 5, 14)
```

```
#   #Loci should have the same no. of 0 as the number of mismatches
#   if (length(Loci$Match[Loci$Match == 0]) != nrow(Mismatch)) {
#     print("Error")
#   } else {
#     return(Loci)
#   }
# }
#
# Analysis.data <- rbind(Mismatch_loci(R154_R1, R154_R2),
#                        Mismatch_loci(R154_R1, R154_R3),
#                        Mismatch_loci(R154_R1, R154_R4),
#                        Mismatch_loci(R155_R1, R155_R2),
#                        Mismatch_loci(R155_R1, R155_R3),
#                        Mismatch_loci(R155_R1, R155_R4),
#                        Mismatch_loci(R156_R1, R156_R2),
#                        Mismatch_loci(R156_R1, R156_R3),
#                        Mismatch_loci(R156_R1, R156_R4)) %>%
#                  mutate(Treatment = factor(Treatment, levels = c("Internal control",
#                                    "Re-used detection plate", "Re-used PCR plate")))
#
# Analysis.data <- left_join(Analysis.data, Nanodrop_SGP, by = "ID")
# Analysis.data$Nanodrop <- as.numeric(Analysis.data$Nanodrop)
#
# n_distinct(Analysis.data$ID) #285 individuals
#
# #We want to exclude the following individuals due to incorrect data
# #"A04_AGP07487" "A05_AGP07488" "A01_AGP16315" "A02_AGP16316" "AGF13004" "AGP16191"
# #(Last two changed name to the original sample)
#
# Analysis.data <- Analysis.data[!Analysis.data$ID %in% c("AGP07487","AGP07488","AGP16315","AGP16316"),]
# n_distinct(Analysis.data$ID) #281 individuals
# write_csv(Analysis.data, file.path(working_dir, "Analysis.data.csv"))
```

To calculate the single-locus genotype error rates, the following function was used to identify and count both mismatched alleles and mismatched single-locus genotypes. It calculates the number of mismatched alleles, mismatched single-locus genotypes and the total number of single-locus genotypes within each comparison between a standard protocol and one of the other treatments, which it uses to calculate an allelic error rate and a single-locus genotyping error rate. It creates the dataset Mismatches.csv, which is later used to calculate per treatment single-locus genotype error rates.

```
# #Function to compare scored alleles between two rounds and find the mismatches
# #The first return statement returns the total mismatches and error rate
#
# Mismatch <- function(DF1, DF2){
#   #Name of the plates
#   Rack_1 <- deparse(substitute(DF1))
#   Rack_2 <- deparse(substitute(DF2))
#
#   if (Rack_1 == "R154_R1") {
#     #On plate R154 the samples in 4th and 5th row are not part of the analysis
#      DF1 <- DF1[-(4:5),]
#      DF2 <- DF2[-(4:5),]
#   } else if (Rack_1 == "R156_R1"){
#       #The repeats are not consistent across all plates, so we remove them
#       DF1 <- DF1[-(1:2),]
#       DF2 <- DF2[-(1:2),]
```

```
#   } else {
#     DF1 <- DF1
#     DF2 <- DF2
#   }
#
#   #To recognize treatment, simplify the 'name' of the DF to the round number (R*)
#   Round <- gsub("R\\d{3}|_", "", Rack_2)
#   if (Round == "R2"){
#     Treatment <- "Internal control"
#   } else if (Round == "R3") {
#     Treatment <- "Re-used detection plate"
#   } else {
#     Treatment <- "Re-used PCR plate"
#   }
#
#   #locate the mismatches by comparing the two data frames and count them
#   mismatches <- which(DF1 != DF2, arr.ind = TRUE)
#   #Some mismatches are caused by NA values, these are excluded
#   #caused by NA
#   na_mismatch <- rbind(subset(mismatches, is.na(DF1[mismatches])),
#                        subset(mismatches, is.na(DF2[mismatches]))) %>%
#                  subset(., !duplicated(.))
#   #remove these false mismatches
#   mismatches <- cbind(mismatches, paste(mismatches[,1],mismatches[,2]))
#   na_mismatch <- cbind(na_mismatch, paste(na_mismatch[,1],na_mismatch[,2]))
#   mismatches <- mismatches[!mismatches[,3] %in% na_mismatch[,3], ]
#   mismatches <- mismatches[,1:2]
#
#   Total_mismatches <- length(mismatches[,1])
#
#   #The allelic error rate is calculated by dividing the number of errors with
#   #the total number of comparisons made between the two dataframes
#   #Without any missing data/errors, this would be 95*78 = 7410
#   #this is 7410/2 = 3705 reactions
#   Total_comparisons <- sum(!is.na(DF1[3:80]) & !is.na(DF2[3:80]))
#   #Allelic error rate
#   Error_rate <- Total_mismatches/Total_comparisons
#
#    #We are also interested in the genotype error rate
#   mismatches_df <- as.data.frame(mismatches)
#   #identify duplicated individuals, by removing unique ID
#   Dup_ID <- mismatches_df %>% group_by(row) %>% filter(n()>1) %>%
#     mutate(row = as.numeric(row),
#            col = as.numeric(col))
#   Dup_ID$combine <- as.numeric(paste(Dup_ID$row, Dup_ID$col, sep = ""))
#   #identify whether the mismatch is in the same locus
#   even <- subset(Dup_ID, Dup_ID$combine %% 2 == 0)
#   uneven <- subset(Dup_ID, Dup_ID$combine %% 2 != 0)
#   #For these the locus is the same
#   Dup_locus <- sum(ifelse((uneven$combine+1) %in% even$combine, 1, 0))
#   #Total number of reactions
#   Total_genotype_errors <- (length(mismatches[,1])-Dup_locus)
#
#   #Genotype error rate
#   Error_rate_genotype <- Total_genotype_errors/(Total_comparisons/2)
```

```
#   No_reactions <- (Total_comparisons/2)
#
#   #The Output is a data frame with the results
#   Output <- as.data.frame(cbind(Rack_1, Rack_2, Treatment,
#                                 Total_mismatches,Total_genotype_errors,
#                                 No_reactions, Error_rate, Error_rate_genotype)) %>%
#     mutate(Total_mismatches = as.numeric(Total_mismatches),
#            Total_genotype_errors = as.numeric(Total_genotype_errors),
#            No_reactions = as.numeric(No_reactions),
#            Error_rate = as.numeric(Error_rate),
#            Error_rate_genotype = as.numeric(Error_rate_genotype))
#   colnames(Output) <- c("Rack 1", "Rack 2", "Treatment",
#                         "No. of mistyped alleles",  "No. of mistyped reactions",
#                         "No. of reactions", "Allelic error rate", "Genotype error rate")
#   return(Output)
# }
#
# #The following applies the mismatch function to the chosen racks and binds the output together
# #The output contains the mismatches and the error rate
# Mismatches <- bind_rows(
#   Mismatch(R154_R1, R154_R2),
#   Mismatch(R154_R1, R154_R3),
#   Mismatch(R154_R1, R154_R4),
#   Mismatch(R155_R1, R155_R2),
#   Mismatch(R155_R1, R155_R3),
#   Mismatch(R155_R1, R155_R4),
#   Mismatch(R156_R1, R156_R2),
#   Mismatch(R156_R1, R156_R3),
#   Mismatch(R156_R1, R156_R4)) %>%
#   arrange(factor(Treatment, levels = c("Internal control",
#                                        "Re-used detection plate",
#                                        "Re-used PCR plate")))
#
# knitr::kable(Mismatches, row.names = F)
# Total_Mcases <- sum(Mismatches$`No. of mismatches`)
# write_csv(Mismatches, file.path(working_dir, "Mismatches.csv"))
```

#### Model

We ran a Bayesian logistic regression mixed model with treatment included as a three level categorical variable to explore the effect on mismatched genotypes data. The 'internal control' treatment was set as the reference category and thereby includes the intercept. Sample ID, DNA plate, multiplexes and loci variables were included as random effects in the model.

```
# #The model takes about 24h to run; model output has been saved as model.mismatch.2.Rdata

# load the data
# Analysis.data <- read.csv(file.path(working_dir, "Analysis.data.csv"))

#A Bernoulli trial is a random experiment that has two possible outcomes: success or failure.
#Bernoulli is a special binary case where the number of trials is 1.

#thin: adjust thinning so you have at least 1000 samples per chain
#(check: (iter-warmup)/thin)
#max_treedepth default = 10 (do not increase beyond 12)
# set.seed(1995)
```

```
# model.mismatch <- brms::brm(Mismatch ~ Treatment + (1|ID) + (1|Rack) + (1|Loci) + (1|Mix),
#               data = Analysis.data,
#               family = bernoulli,
#               cores = 3,
#               iter = 100000,
#               thin = 70,
#               warmup = 30000, #standard is 20-50% of iter
#               chains = 3,
#               control = list(adapt_delta = 0.99, max_treedepth = 11),
#               backend = "cmdstanr", #improve speed
#               silent = 0) #to see progress
#
# save(model.mismatch, file = "Data/Working_data/Processed_data/model.mismatch.2.Rdata")
```

##### Model diagnostics

We explore the model diagnostics from the brms package. Plot is used to visually investigate the chains and posterior distributions and pp_check (posterior predictive checking) is used to compare the observed outcome variable to a simulated dataset from the posterior predictive distribution.

```
#Load the model information to perform model diagnostics
load(file="Data/Working_data/Processed_data/model.mismatch.2.Rdata")

#Check model
#Trace plots: explore the trace plots of the chains.
#They should be harmonized and bounce up and down.
#Fixed effects
plot(model.mismatch, variable = c("b_Intercept",
                                  "b_TreatmentReMuseddetectionplate",
                                  "b_TreatmentReMusedPCRplate"))
```

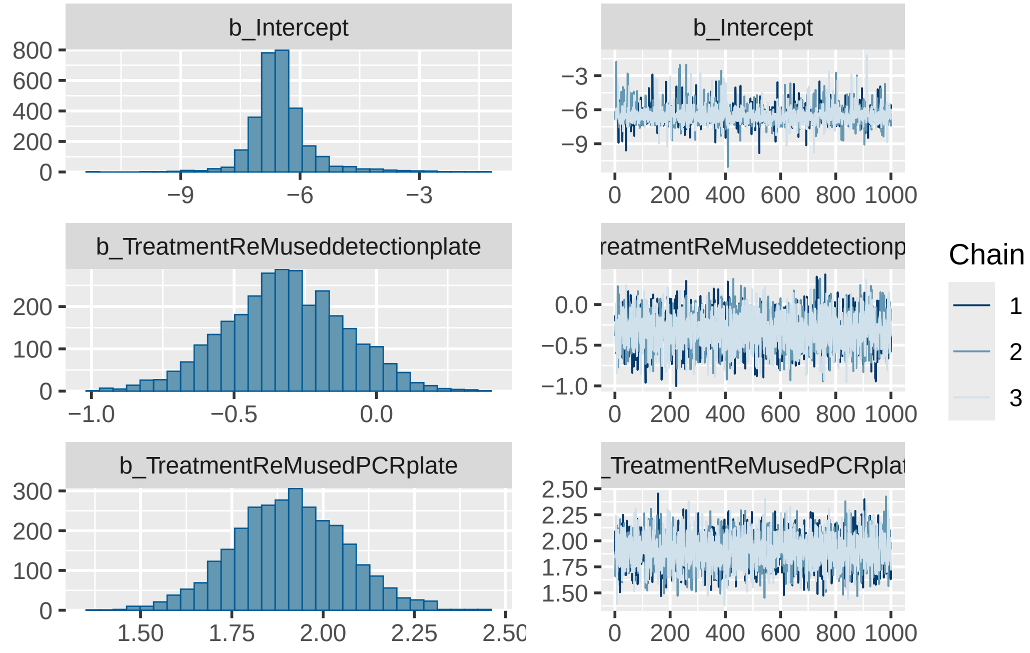

```r
#Random effects
plot(model.mismatch, variable = c("sd_ID__Intercept", "sd_Loci__Intercept",
                                  "sd_Mix__Intercept", "sd_Rack__Intercept"))
```

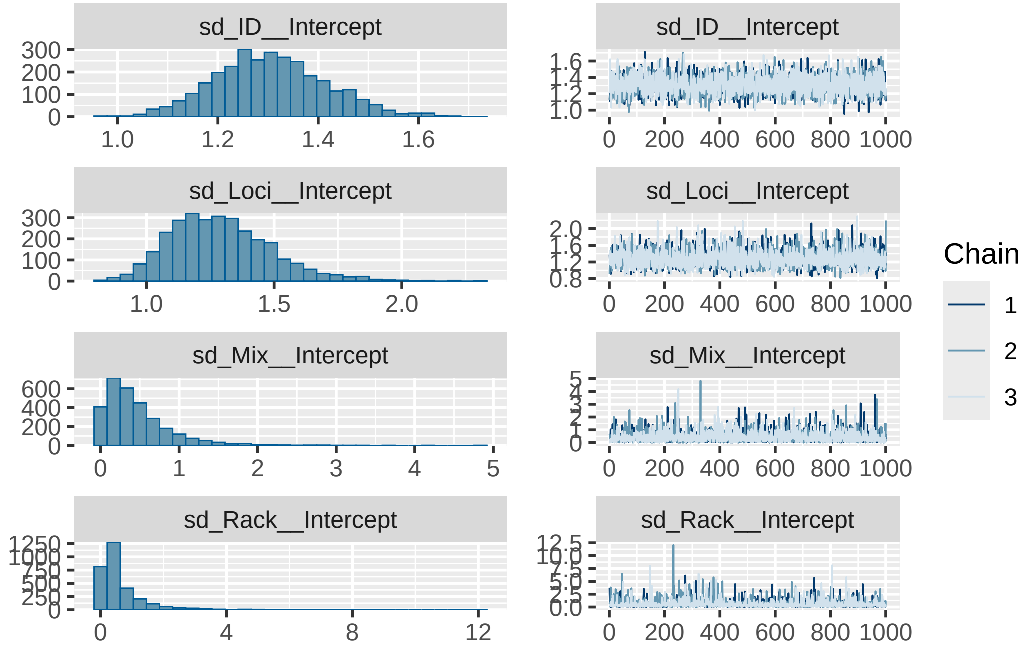

```r
#Further tests
#The 'acf' is an autocorrelation plot used to check for randomness in the dataset.
#if the autocorrelation value is close to 0, randomness can be assumed.
mcmc_plot(model.mismatch, type = "acf") #autocorrelation is close to 0
```

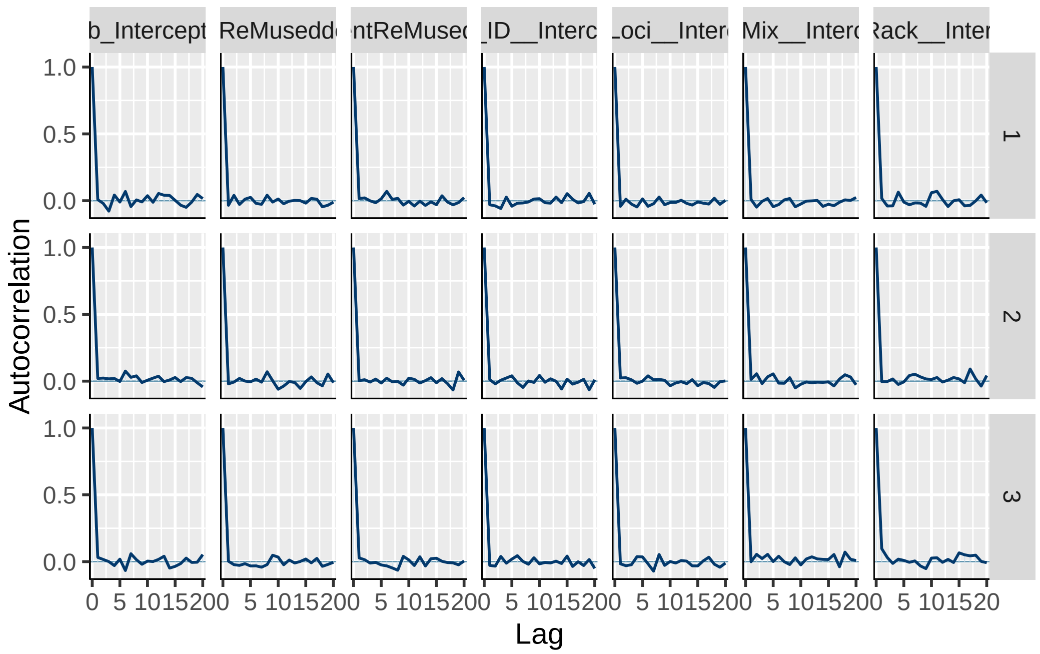

```r
pp_check(model.mismatch, type = "bars", ndraws = 1000)
```

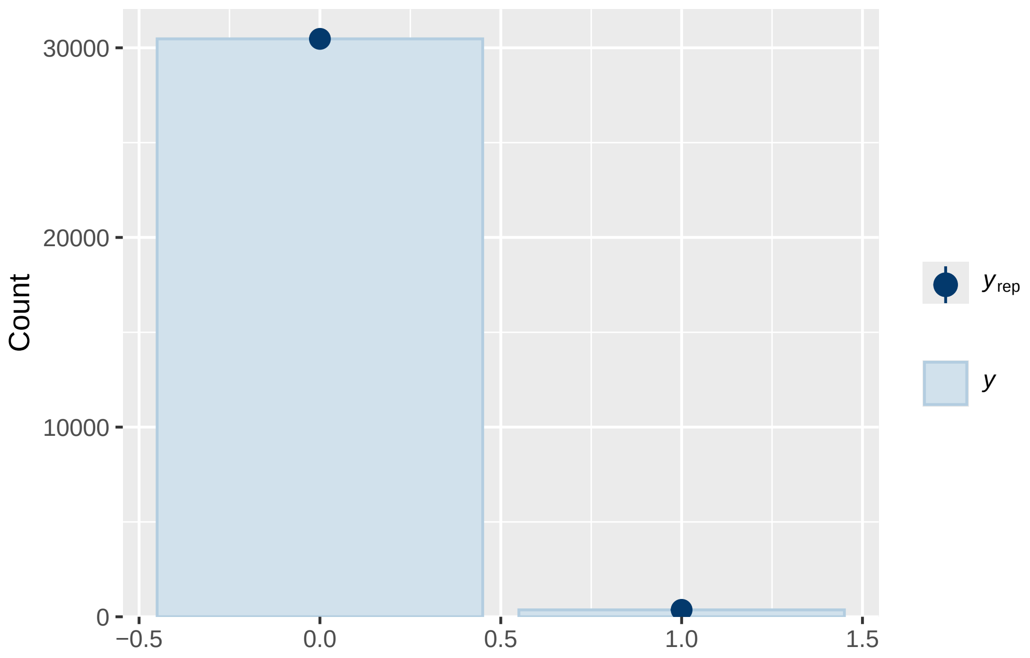

**Rhat**

Rhat statistics are used to diagnose sampling behaviour and should be close to 1. Rhat values above 1 means the models has not yet converged.

```r
#rhat
mcmc_rhat(brms::rhat(model.mismatch))
```

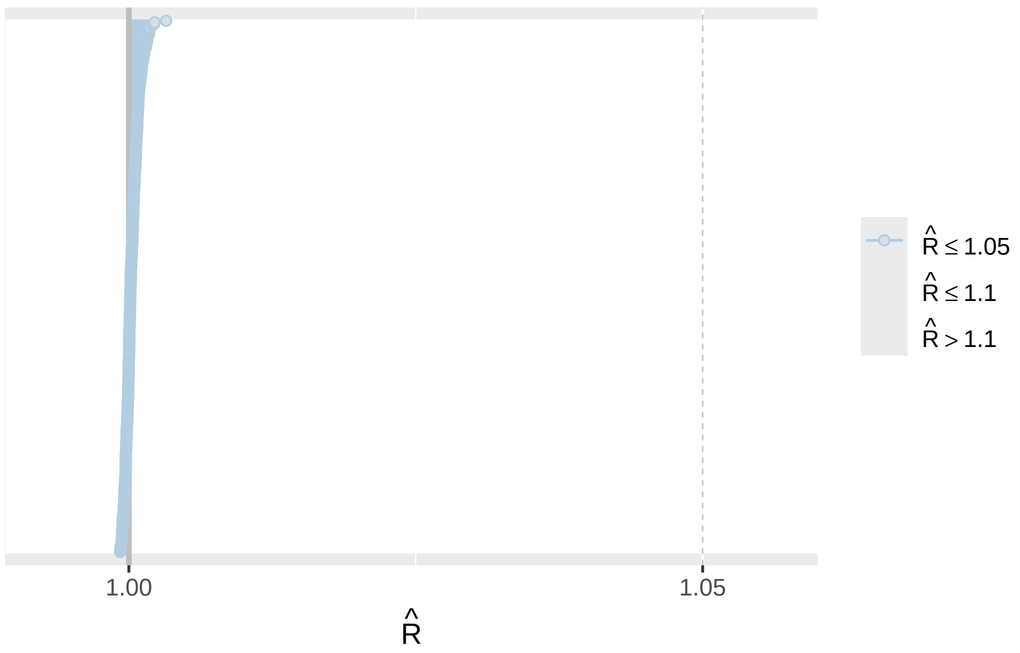

##### Pairs plot

The pairs plot is used to identify collinearity between variables.

```
#To make the pairs plot fit the pdf, we save it and reload it back in.
#png(filename = file.path(figs_dir, "pairs.plot.png"), width = 15,
#                          height = 15, units = "in", res = 300)
#mcmc_plot(model.mismatch, type = "pairs")
#dev.off()
```

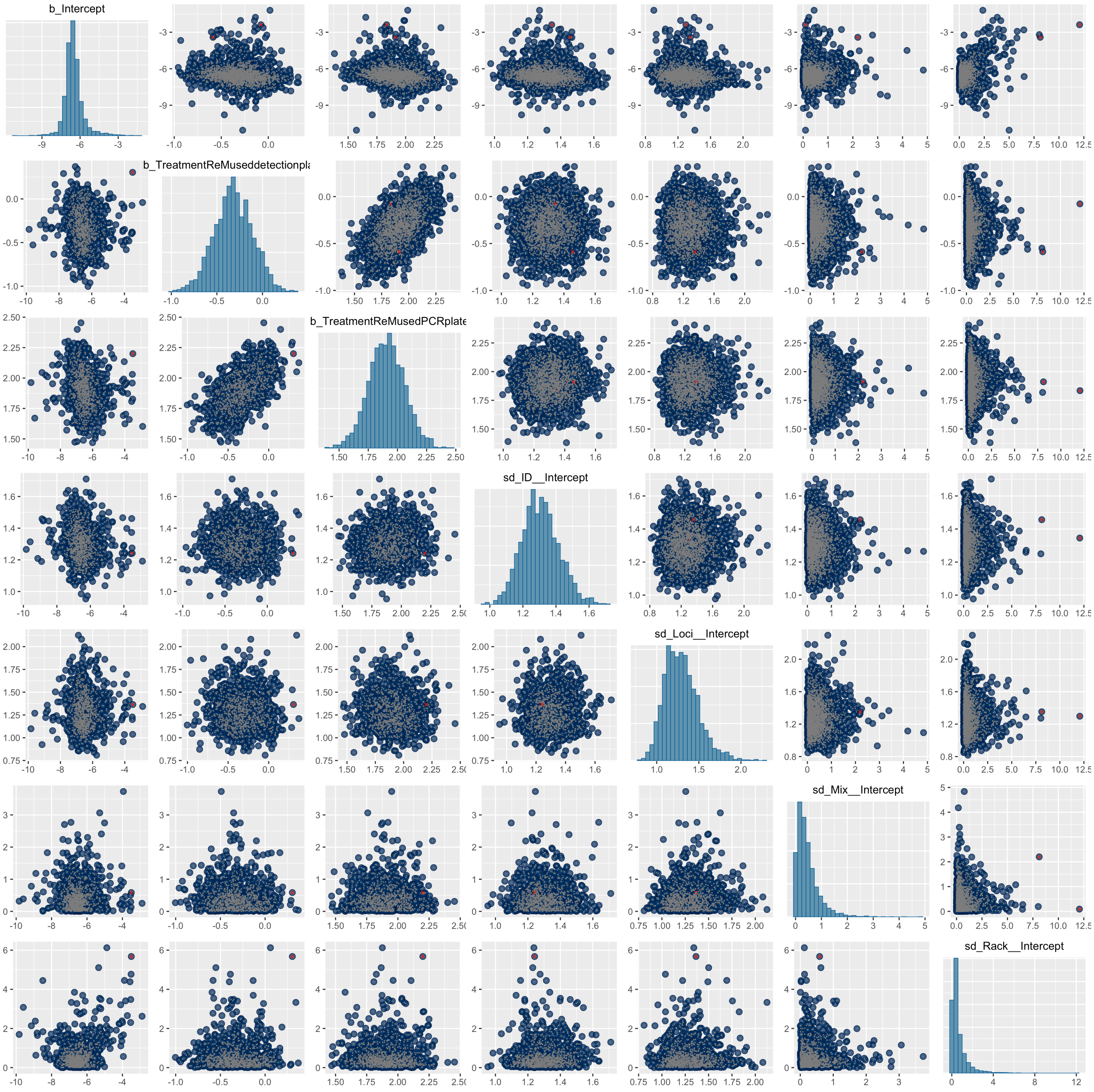

Figure 2: Pairs plot

##### Model summary

The model summary provides general information about the model, group and population level effects. In addition, it provides information on how well the model could estimate the posterior distribution. ESS-bulk and ESS_tail values should be close to 3000 (the total post-warmup draws).

```
#Check model output
#
#The 'internal control' treatment is included in the intercept
#The other treatment groups are 'in reference' to the internal control treatment.
summary(model.mismatch)
```

```
Warning: There were 3 divergent transitions after warmup. Increasing
adapt_delta above 0.99 may help. See
http://mc-stan.org/misc/warnings.html#divergent-transitions-after-warmup
```

```
 Family: bernoulli
  Links: mu = logit
Formula: Mismatch ~ Treatment + (1 | ID) + (1 | Rack) + (1 | Loci) + (1 | Mix)
   Data: Analysis.data (Number of observations: 30835)
  Draws: 3 chains, each with iter = 1e+05; warmup = 30000; thin = 70;
         total post-warmup draws = 3000

Multilevel Hyperparameters:
~ID (Number of levels: 280)
              Estimate Est.Error l-95% CI u-95% CI Rhat Bulk_ESS Tail_ESS
sd(Intercept)     1.31      0.11     1.10     1.53 1.00     3083     2887

~Loci (Number of levels: 39)
              Estimate Est.Error l-95% CI u-95% CI Rhat Bulk_ESS Tail_ESS
sd(Intercept)     1.29      0.20     0.97     1.76 1.00     3089     2778

~Mix (Number of levels: 5)
              Estimate Est.Error l-95% CI u-95% CI Rhat Bulk_ESS Tail_ESS
sd(Intercept)     0.46      0.43     0.02     1.57 1.00     2634     2988

~Rack (Number of levels: 3)
              Estimate Est.Error l-95% CI u-95% CI Rhat Bulk_ESS Tail_ESS
sd(Intercept)     0.64      0.80     0.02     2.86 1.00     2751     2864

Regression Coefficients:
                               Estimate Est.Error l-95% CI u-95% CI Rhat
Intercept                         -6.49      0.75    -7.66    -4.57 1.00
TreatmentReMuseddetectionplate    -0.32      0.21    -0.74     0.08 1.00
TreatmentReMusedPCRplate           1.91      0.15     1.60     2.21 1.00
                               Bulk_ESS Tail_ESS
Intercept                          2984     2788
TreatmentReMuseddetectionplate     3104     3122
TreatmentReMusedPCRplate           2881     2702

Draws were sampled using sample(hmc). For each parameter, Bulk_ESS
and Tail_ESS are effective sample size measures, and Rhat is the potential
scale reduction factor on split chains (at convergence, Rhat = 1).
```

### Results

281 samples were included in the analysis. On rack 154 the two samples from 2007-2008 were excluded, due to lack of DNA for the last treatment. Likewise for two of the positive controls on R156. This resulted in a total of 30.835 observations.

#### Tables

##### Missing data

Percentage of per-reaction missing data for the internal control, reused detection plate and reused PCR plate treatment groups.

```
#Table containing the missing data rate for each treatment
#SLG = single locus genotypes

Missing <- rbind(nrow(filter(Analysis.data, Treatment == "Internal control" & Missing == "0")),
                 nrow(filter(Analysis.data, Treatment == "Re-used detection plate" & Missing == "0")),
                 nrow(filter(Analysis.data, Treatment == "Re-used PCR plate" & Missing == "0")))

SLG <- rbind(nrow(filter(Analysis.data, Treatment == "Internal control")),
             nrow(filter(Analysis.data, Treatment == "Re-used detection plate")),
             nrow(filter(Analysis.data, Treatment == "Re-used PCR plate")))

Missing_rate <- rbind(round((Missing[1]/SLG[1])*100,3),
                      round((Missing[2]/SLG[2])*100,3),
                      round((Missing[3]/SLG[3])*100,3))
Treatment <- rbind("Internal control", "Reused detection plate", "Reused PCR plate")

Table_missing_rate <- as.data.frame(cbind(Treatment, SLG, Missing, Missing_rate))
Table_missing_rate <- Table_missing_rate %>%
  mutate(., V2 = as.numeric(V2), V3 = as.numeric(V3), V4 = as.numeric(V4))

colnames(Table_missing_rate) <- c("Treatment", "No. of single-locus genotypes",
                                  "No. of missing data", "Missing data %")

Table_missing_rate <- gt(Table_missing_rate)
Table_missing_rate <- Table_missing_rate %>%
  fmt_number(columns = 2:3, decimals = 0, sep_mark = ".", dec_mark = ",") %>%
  fmt_number(columns = 4, decimals = 3, sep_mark = ".", dec_mark = ",")

Table_missing_rate %>%
  tab_style(style = cell_text(color = 'black', weight = "bold"),
            locations = cells_body(columns = 1)) %>%
  tab_options(table.font.names = "TNR",column_labels.font.weight = "bold", table.width = 700) %>%
  cols_align(align = "center", columns = c(2,3,4)) %>%
  tab_style(style = cell_fill(color = 'grey90'),
            locations = cells_body(rows = c(1,3))) -> Table.missing.finished

#library(webshot2)
#gtsave(Table.missing.finished, file.path(tabs_dir, "Missingrates.table.png"))
```

Table 2: Missing data

| Treatment | No. of single-locus genotypes | No. of missing data | Missing data (%) |
|---|---|---|---|
| Internal control | 10.959 | 322 | 2,938 |
| Reused detection plate | 10.959 | 248 | 2,263 |
| Reused PCR plate | 10.959 | 1.472 | 13,432 |

##### Genotyping error rates

Per-reaction genotyping error rates for the internal control, reused detection plate and reused PCR plate treatment groups, calculated relative to the standard protocol treatment group.

```
#load data
Mismatches <- read.csv(file.path(working_dir, "Mismatches.csv"))

#Table containing the error rates for each treatment
Errors <- rbind(sum(Mismatches$`No..of.mistyped.reactions`[1:3]),
                sum(Mismatches$`No..of.mistyped.reactions`[4:6]),
                sum(Mismatches$`No..of.mistyped.reactions`[7:8]))
Reactions <- rbind(sum(Mismatches$`No..of.reactions`[1:3]),
                   sum(Mismatches$`No..of.reactions`[4:6]),
                   sum(Mismatches$`No..of.reactions`[7:8]))
Error_rate <- rbind(round(Errors[1]/Reactions[1],3),
                    round(Errors[2]/Reactions[2],3),
                    round(Errors[3]/Reactions[3],3))
Treatment <- rbind("Internal control", "Reused detection plate", "Reused PCR plate")

Table_error_rate <- as.data.frame(cbind(Treatment, Reactions, Errors, Error_rate))
Table_error_rate$V2 <- as.numeric(Table_error_rate$V2)
colnames(Table_error_rate) <- c("Treatment",
                                "No. of single-locus genotypes",
                                "No. of mismatches",
                                "Genotype error rate")

Table_error_rate <- gt(Table_error_rate)
Table_error_rate <- Table_error_rate %>%
  fmt_number(columns = 2, decimals = 0, sep_mark = ".", dec_mark = ",")

Table_error_rate %>%
  tab_style(style = cell_text(color = 'black', weight = "bold"),
            locations = cells_body(columns = 1)) %>%
  tab_options(table.font.names = "TNR",column_labels.font.weight = "bold", table.width = 700) %>%
  cols_align(align = "center", columns = c(2,3,4)) %>%
  tab_style(style = cell_fill(color = 'grey90'),
            locations = cells_body(rows = c(1,3))) -> Table.error.finished

#save table as png
#library(webshot2)
#gtsave(Table.error.finished, file.path(tabs_dir, "Errorrates.table.png"))
```

Table 3: Genotyping error rate

| Treatment | No. of single-locus genotypes | No. of mismatches | Genotyping error rate |
|---|---|---|---|
| Internal control | 10.541 | 57 | 0,005 |
| Reused detection plate | 10.599 | 42 | 0,004 |
| Reused PCR plate | 6.159 | 210 | 0,034 |

##### Model output

Point beta estimates and 95% confidence intervals from the Bayesian logistic mixed effect model testing for the effects of the fixed effect "treatment" on genotype errors while controlling for the random effects sample ID, DNA plate, multiplexed reaction and loci.

```
#make a table for the model output
models <- list("Genotype errors" = model.mismatch)

#estimates stem from model summary
rows <- tribble(~term, ~`Genotype errors`,
                'Fixed effects', '',
                'Random effects', '')
attr(rows, 'position') <- c(1,5)

#order the values in the table
cm <- c('b_Intercept'    = 'Internal control',
        'b_TreatmentReMuseddetectionplate' = 'Reused detection plate',
        'b_TreatmentReMusedPCRplate'    = 'Reused PCR plate',
        'sd_ID__Intercept' = 'Sample ID', 'sd_Rack__Intercept' = 'DNA plate',
        'sd_Mix__Intercept' = 'Multiplexed reaction', 'sd_Loci__Intercept' = 'Locus')

f <- function(x) format(round(x, 3), big.mark=",", decimal.mark=".")
gm <- list(list("raw" = "nobs", "clean" = "Number of observations", "fmt" = f),
           list("raw" = "r.squared", "clean" = "R²", "fmt" = f),
           list("raw" = "r2.marginal", "clean" = "Marginal R²", "fmt" = f))

Table <- modelsummary::modelsummary(models,
                           estimate  = "{estimate} [{conf.low}, {conf.high}]", statistic = NULL,
                           conf_level = .95, metrics = "R2", add_rows = rows,
                           fmt = f, coef_map = cm, gof_map = gm, output = "gt")

Table %>%
  tab_style(style = cell_text(color = 'black', weight = "bold"),
            locations = cells_body(columns = 1)) %>%
  tab_options(table.font.names = "TNR",column_labels.font.weight = "bold", table.width = 400) %>%
  tab_style(style = cell_fill(color = 'grey95'),
            locations = cells_body(rows = c(3,7,9,11))) %>%
  tab_style(style = cell_fill(color = 'grey90'),
            locations = cells_body(rows = c(1,5))) -> Table.finished

#library(webshot2)
#gtsave(Table.finished, file.path(tabs_dir, "Model.output.errorrates.png"))
```

Table 4: Model output

| | Beta estimates |
|---|---|
| Fixed effects | |
| Internal control | -6,572 [-7,658, -4,565] |
| Reused detection plate | -0,320 [-0,739, 0,083] |
| Reused PCR plate | 1,907 [1,602, 2,209] |
| Random effects | |
| Sample ID | 1,302 [1,100, 1,533] |
| DNA plate | 0,388 [0,025, 2,864] |
| Multiplexed reaction | 0,348 [0,016, 1,571] |
| Locus | 1,273 [0,970, 1,759] |
| Number of observations | 30.835 |
| $R^2$ | 0,117 |
| Marginal $R^2$ | 0,001 |

#### Figures

##### Posterior beta estimates

Posterior distributions of the beta coefficients of the internal control (blue), reused detection plate (orange) and reused PCR plate (red) treatment groups on genotyping errors. Point beta estimate represented with a grey dot. 50% and 90% confidence intervals are represented with thick and thin black lines respectively. Reused detection plate and reused PCR plate treatment group values has been added/subtracted from the intercept to visually represent the 'actuaæ' beta estimates for each group instead of the relative beta estimate to the intercept group 'internal control'.

```
# Plot the model
#Extract info from model output
model_interval_mismatch <- mcmc_intervals_data(model.mismatch)
model_interval_mismatch <- subset(model_interval_mismatch, grepl("b_", parameter)) %>%
  select(-outer_width, -inner_width, -point_est)

model_interval_mismatch <- model_interval_mismatch %>%
  mutate(parameter = case_when(
  parameter == "b_Intercept"  ~ "Internal control",
  parameter == "b_TreatmentReMuseddetectionplate" ~ "Reused detection plate",
  parameter == "b_TreatmentReMusedPCRplate"  ~ "Reused PCR plate"))

#M value for intercept
m_mismatch <- model_interval_mismatch %>%
  subset(parameter == "Internal control") %>%
  select(m)

#Add intercept values to estimates
Intercept_mismatch <- model_interval_mismatch %>%
  subset(parameter == "Internal control")
Other_mismatch <- model_interval_mismatch  %>%
  subset(parameter != "Internal control") %>%
  mutate(.[2:6]+m_mismatch$m)
model_interval_mismatch <- rbind(Intercept_mismatch, Other_mismatch)

#area (for density clouds)
model_info_mismatch <- mcmc_areas_data(model.mismatch)
#model_info <- subset(model_info, !grepl("b_intercept", parameter))
model_info_mismatch <- subset(model_info_mismatch, grepl("b_", parameter))
model_info_mismatch <- model_info_mismatch %>% mutate(parameter = case_when(
  parameter == "b_Intercept"  ~ "Internal control",
  parameter == "b_TreatmentReMuseddetectionplate" ~ "Reused detection plate",
  parameter == "b_TreatmentReMusedPCRplate"  ~ "Reused PCR plate"))

model_info_mismatch$parameter <- factor(model_info_mismatch$parameter,
                                        levels = c("Internal control",
                                                   "Reused detection plate",
                                                   "Reused PCR plate"))
#add median value to other treatments
model_info_mismatch_intercept <- model_info_mismatch %>%
  subset(parameter == "Internal control")
model_info_mismatch_other <- model_info_mismatch %>%
  subset(parameter != "Internal control") %>%
  mutate(x = x+m_mismatch$m)
model_info_mismatch <- rbind(model_info_mismatch_intercept, model_info_mismatch_other)
```

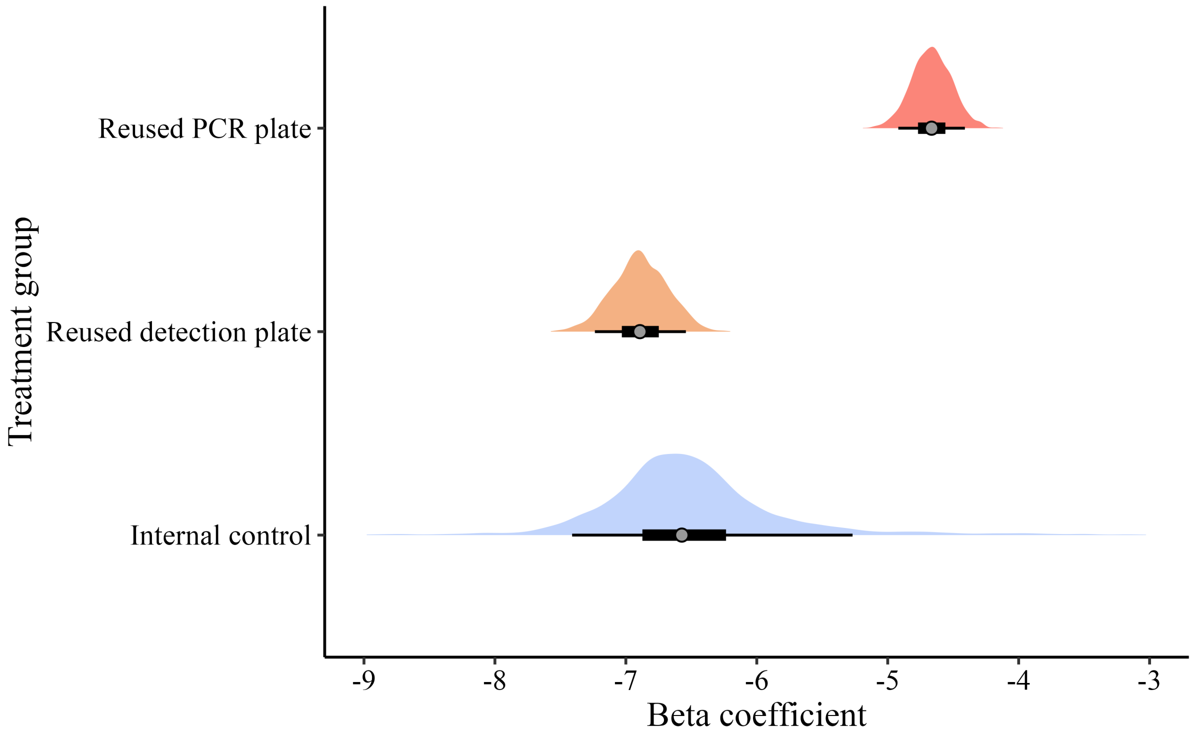

Figure 3: Model: posterior beta estimates

#### Session information

```
R version 4.2.1 (2022-06-23)
Platform: x86_64-apple-darwin17.0 (64-bit)
Running under: macOS Big Sur ... 10.16

Matrix products: default
BLAS:   /Library/Frameworks/R.framework/Versions/4.2/Resources/lib/libRblas.0.dylib
LAPACK: /Library/Frameworks/R.framework/Versions/4.2/Resources/lib/libRlapack.dylib

locale:
[1] en_US.UTF-8/en_US.UTF-8/en_US.UTF-8/C/en_US.UTF-8/en_US.UTF-8

attached base packages:
[1] stats     graphics  grDevices utils     datasets  methods   base

other attached packages:
 [1] webshot2_0.1.1     gt_0.11.0          ggridges_0.5.4     bayesplot_1.9.0
 [5] brms_2.22.0        Rcpp_1.0.11        rstan_2.32.5       StanHeaders_2.32.5
 [9] kableExtra_1.4.0   rstatix_0.7.0      knitr_1.45         png_0.1-7
[13] stringi_1.8.3      here_1.0.1         lubridate_1.9.2    forcats_1.0.0
[17] stringr_1.5.1      purrr_1.0.2        readr_2.1.4        tidyr_1.3.0
[21] tibble_3.2.1       ggplot2_3.5.1      tidyverse_2.0.0    dplyr_1.1.3

loaded via a namespace (and not attached):
  [1] websocket_1.4.2      TH.data_1.1-1        colorspace_2.1-0
  [4] ellipsis_0.3.2       rprojroot_2.0.3      estimability_1.4.1
  [7] QuickJSR_1.1.3       parameters_0.22.0    rstudioapi_0.15.0
 [10] listenv_0.9.0        farver_2.1.1         chromote_0.3.1
```

```
 [13] fansi_1.0.6          mvtnorm_1.2-2        xml2_1.3.3
 [16] bridgesampling_1.1-2 codetools_0.2-18     splines_4.2.1
 [19] jsonlite_1.8.9       broom_1.0.5          effectsize_0.8.9
 [22] modelsummary_2.1.1   compiler_4.2.1       emmeans_1.8.7
 [25] backports_1.5.0      Matrix_1.5-3         fastmap_1.1.1
 [28] cli_3.6.3            later_1.3.0          htmltools_0.5.8.1
 [31] prettyunits_1.1.1    tools_4.2.1          coda_0.19-4
 [34] gtable_0.3.4         glue_1.8.0           reshape2_1.4.4
 [37] posterior_1.6.0      tables_0.9.17        V8_4.2.1
 [40] carData_3.0-5        vctrs_0.6.5          svglite_2.1.2
 [43] nlme_3.1-163         insight_0.20.1       tensorA_0.36.2.1
 [46] xfun_0.41            globals_0.16.2       ps_1.8.0
 [49] timechange_0.2.0     lifecycle_1.0.4      pacman_0.5.1
 [52] future_1.33.0        MASS_7.3-57          zoo_1.8-11
 [55] scales_1.3.0         ragg_1.2.5           hms_1.1.2
 [58] promises_1.2.0.1     Brobdingnag_1.2-9    parallel_4.2.1
 [61] sandwich_3.0-2       inline_0.3.19        yaml_2.3.8
 [64] curl_4.3.3           gridExtra_2.3        loo_2.8.0
 [67] bayestestR_0.13.2    checkmate_2.3.2      pkgbuild_1.3.1
 [70] cmdstanr_0.8.1.9000  rlang_1.1.4          pkgconfig_2.0.3
 [73] systemfonts_1.0.5    matrixStats_1.4.1    distributional_0.5.0
 [76] evaluate_0.23        lattice_0.20-45      labeling_0.4.3
 [79] rstantools_2.2.0     processx_3.8.4       tidyselect_1.2.0
 [82] parallelly_1.36.0    plyr_1.8.7           magrittr_2.0.3
 [85] R6_2.5.1             generics_0.1.3       multcomp_1.4-20
 [88] pillar_1.9.0         withr_3.0.1          datawizard_0.11.0
 [91] survival_3.3-1       abind_1.4-8          performance_0.12.0
 [94] future.apply_1.11.2  crayon_1.5.2         car_3.1-1
 [97] utf8_1.2.4           tzdb_0.3.0           rmarkdown_2.25
[100] grid_4.2.1           data.table_1.16.0    callr_3.7.3
[103] digest_0.6.33        xtable_1.8-4         textshaping_0.3.6
[106] RcppParallel_5.1.5   stats4_4.2.1         munsell_0.5.0
[109] viridisLite_0.4.2
```
